# Generation of CRISPR-edited birch Plants without DNA integration using Agrobacterium-mediated Transformation Technology

**DOI:** 10.1101/2023.10.09.561573

**Authors:** Shilin Sun, Xue Han, Ruoxuan Jin, Junbo Jiao, Jingwen Wang, Siyuan Niu, Ziyao Yang, Di Wu, Yucheng Wang

**Affiliations:** College of Forestry, Shenyang Agricultural University, Shenyang 110866, China; Key Laboratory of Forest Tree Genetics, Breeding and Cultivation of Liaoning Province, Shenyang 110866, China

**Author notes:** Corresponding author: Yucheng Wang College of Forestry, Shenyang Agricultural University, Shenyang 110866, China. These authors contributed equally to this work (Shilin Sun, Xue Han).

## Abstract

CRISPR/Cas9 system has emerged as a powerful tool in genome editing; however, generation of CRISPR-edited DNA-free plants is still challenging. In this study, *Betula platyphylla* (birch) was used to build a method to generate CRISPR-edited plant without foreign DNA integration using Agrobacterium-mediated transformation (CPDAT method). This technique utilizes transient genetic transformation to introduce T-DNA coding gRNA and Cas9 into birch cells, and T-DNA will express to synthesize gRNA and Cas9 protein, which will form a complex to cleave the target DNA site. The genome may be mutated due to DNA repair, and these mutations will be preserved and accumulated not dependent on whether T-DNA is integrated into the genome or not. After transient transformation, birch plants were cut into explants to induce adventitious buds without antibiotic selection pressure. Each adventitious bud can be considered as an independent potentially CRISPR-edited line for mutation detection. CRISPR-edited birch plants without foreign DNA integration are further selected by screening CRISPR-edited lines without T-DNA integration. Among 65 randomly chosen independent lines, the mutation rate was 80.00% including 40.00% of lines with both alleles mutated. In addition, 5 lines out of 65 studied lines (7.69%) were CRISPR-edited birch plants without DNA integration. In conclusion, this innovative method presents a novel strategy for generating CRISPR-edited birch plants, thereby significantly enhancing the efficiency of generating common CRISPR-edited plants. These findings offer considerable potential to develop plant genome editing techniques further.

## Introduction

CRISPR/Cas9 is a robust and effective gene-editing technology for plant breeding, which can precisely and efficiently modify a genome(Fidan et al., 2023). CRISPR/Cas system was initially found to recognize and cleave the invading viruses or phages’ DNA serving as an immune system in bacterium(Ahmad, 2023; Kim et al., 2016; Komor et al., 2016; Koonin et al., 2017; Zetsche et al., 2015). CRISPR-Cas systems have been classified into two main classes (Class 1 and 2) and various types (I, II, III, IV, V, and VI) based on their CRISPR-Cas loci arrangement and associated Cas protein(Koonin et al., 2017; Makarova and Koonin, 2015). The two main classes are class 1 and 2 which are distinguished according to their utilized complex of CRISPR RNA (crRNA)-effector protein(McDonald et al., 2019). Class 1 systems (including type I, III, and IV) are constituted by the mature crRNA bound by several Cas proteins to form a huge protein complex. This complex serves as a guide to the target DNA site by pairing between the complementary strand DNA of the protospacer (target site) and the 5’-end sequence of the gRNA spacer and also has nuclease activity for cleaving the targeted sequences(Garneau et al., 2010; Tiwari et al., 2023; Wiedenheft et al., 2011). The class 2 systems include type II, V, and VI, which has only one Cas protein, such as Cas9, Cas12, or Cas13 respectively, and also serve the function of targeting and cleaving DNA(Jinek et al., 2012). Among class 2 systems, the Cas9-CRISPR system has been widely applied.

In this system, a single-guide RNA (sgRNA) is formed by merging trans-activating CRISPR RNA (tracrRNA) and CRISPR-RNA (crRNA) to create a loop at the duplex region end. Streptococcus pyogenes Cas9 (SpCas9) is the most commonly used Cas protein, which recognizes the DNA target sequence requiring a short protospacer-adjacent motif (PAM) with the sequence of 5′-NGG-3′(Legut et al., 2020). The sgRNA binds to Cas9 and guides the Cas9 protein to induce site-specific DNA double-strand breaks (DSBs)(Goyon et al., 2021). In this process, the cells may employ two main mechanisms to repair DSBs, i.e., non-homologous end joining (NHEJ) or homology-directed repair (HDR). The NHEJ is an error-prone mechanism, which enables substitutions, random small deletions, or insertions, and probably leads to gene knock-out (KO) mutations. The HDR mechanism usually generates point deletions or mutations caused by gene knock-in (KI)(Schrank et al., 2018; Ye et al., 2018).

Traditionally, CRISPR editing in plants has been done by using a stable transformation, integrating Cas protein and single guide RNA (sgRNA) into the plant genome(Metje-Sprink et al., 2019). However, these transformation methods have the weakness that had introduced foreign DNA and integrating it into the genome, which also induces concerns about genetically modified organisms (GMOs) as transgenic plants. As an alternative approach, a DNA-free CRISPR system employed plasmids harboring the sgRNA and Cas sequences or CRISPR ribonucleoprotein (RNP) that can be delivered directly into protoplasts using transient genetic transformation or delivered pre-assembled CRISPR/Cas9 ribonucleoproteins (RNPs) into plant cells by using particle bombardment, allowing recombinant DNA-free plants to be generated(Andersson et al., 2018; De Bruyn et al., 2020; Hsu et al., 2019; Hsu et al., 2020; Liang et al., 2017; Liang et al., 2018; Lin et al., 2022; Woo et al., 2015). These technologies are completely free of DNA so the problem of integration of DNA into the genome has been excluded. These technologies are of great importance for the reason that they represent an innovative non-genetically modified organisms (GMOs) approach that can avoid the GMO debate. At the same time, the generation of CRISPR-edited plants without DNA integration is mainly using transient transformation to introduce plasmid into protoplasts and using particle bombardment to introduce pre-assembled CRISPR/Cas9 RNPs. These methods are involved in preparing protoplasts or pre-assembled CRISPR/Cas9 RNPs, protoplast differentiated into buds. Despite the advances, these methods remain complex and time-consuming. Therefore, a more simple and efficient method needs to be developed.

Genetic transformation mediated by *Agrobacterium tumefaciens* is a commonly used technology for plant CRISPR genome editing up to now(Aliu et al., 2022; Lopez-Casado et al., 2023; Tang et al., 2023). However, there is no report about using this method to generate CRISPR-edited plants without DNA integration, and an increase in the efficiency of CRISPR transformation mediated by *A. tumefaciens* is still an important problem that limits plant genome editing.

In the present study, we introduce a novel approach that employs genetic transformation mediated by *A. tumefaciens* to generate CRISPR-edited plants. This method can improve transformation efficiency substantially compared with the classic gene transfer mediated by *A. tumefaciens*, and can also be used to efficiently generate genome editing plants without foreign DNA integration. Therefore, this method may have a wide application in genome editing mediated by CRISPR/Cas.

## Results

### The principle of Generating CRIPSR-edited plants without DNA-insertion using the Agrobacterium-mediated transformation method (CPDAT)

A method for the generation of CRISPR-edited birch plants without DNA integ ration was built. In this method based on Agrobacterium-mediated transformation, the T-DNA coding gRNA and Cas9 were introduced into the cell nucleus using the transient transformation method, then gRNA and Cas9 are expressed an d form a complex to cleave the target DNA site on the genome, which may induce DNA mutation during DNA repair with NHEJ mechanism. The mutation on the genome will be preserved in a genome that is not dependent on whether T-DNA integration or not. Additionally, mutations in one cell or several cells might accumulate, increasing the frequency of the formation of adventitious buds with mutated genomes. This is the reason why two rounds of heat treatment are performed which can increase the quantity of cleaving genome leading to increased DNA mutation. Therefore, when the cells with genome mutation a re differentiated into adventitious buds, the genome mutation will also be passed on to the cells that are differentiated into adventitious buds (Fig. 1). This method can generate CRISPR-edited birch plants efficiently, and CRISPR-edited plants without DNA integration can be further selected from the CRISPR-edited plants by detecting whether T-DNA integrated into genome or not.

**Fig 1.**
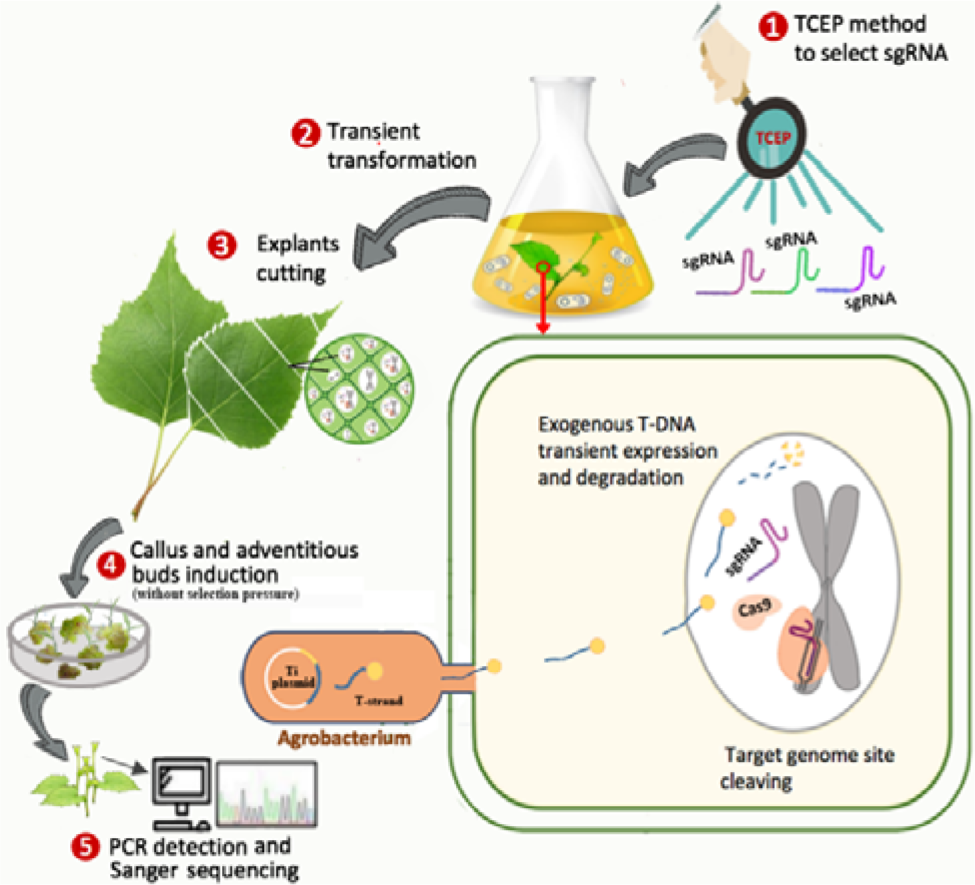
Principle of the CPDAT method. The T-DNA containing gRNA and Cas9 are delivered into plant cells using the transient transformation method, and the target genome site will be cleaved to induce mutation, which is not dependent on whether T-DNA integration or not. The mutation in cells will be accumulated and will be passed on to the cells and the cells then differentiate into adventitious buds. The CRISPR-edited pla nts including CRISPR-edited plants without DNA integration can be identified by Cruiser^TM^ digestion, Sanger sequencing, and PCR detection.

### The procedure of CPDAT

The procedure for generating CRISPR-edited birch plants without DNA integration was as the following. Step 1, select the gRNA site that can be cleaved effectively by Cas9 using the TCEP method. Step 2, transient transformation is performed on the entire plant, which enables T-DNA containing gRNA and Cas9 to be delivered into birch plant cells to cleave the target DNA site. In step 3, heat (37°C) treatment is used to enhance Cas9 activity on cleaving the target site; in this step, the cleavage efficiency should be determined using TCEP after each heat treatment. Step 4: The transiently transformed plants are separated into explants without antimicrobial selection pressure to promote adventitious b uds. Step 5, the adventitious buds grow into small plantlets. Step 6, determination of the mutation induced by CRISPR. Step 7 involves screening CRISPR-edited plants without DNA integration using PCR to determine whether or not T -DNA integration has occurred (Fig. 2). If the generation of common CRISPR-edited plants (i.e., not CRISPR-edited plants without DNA integration) is needed, only steps 1 to 5 are needed.

**Fig 2.**
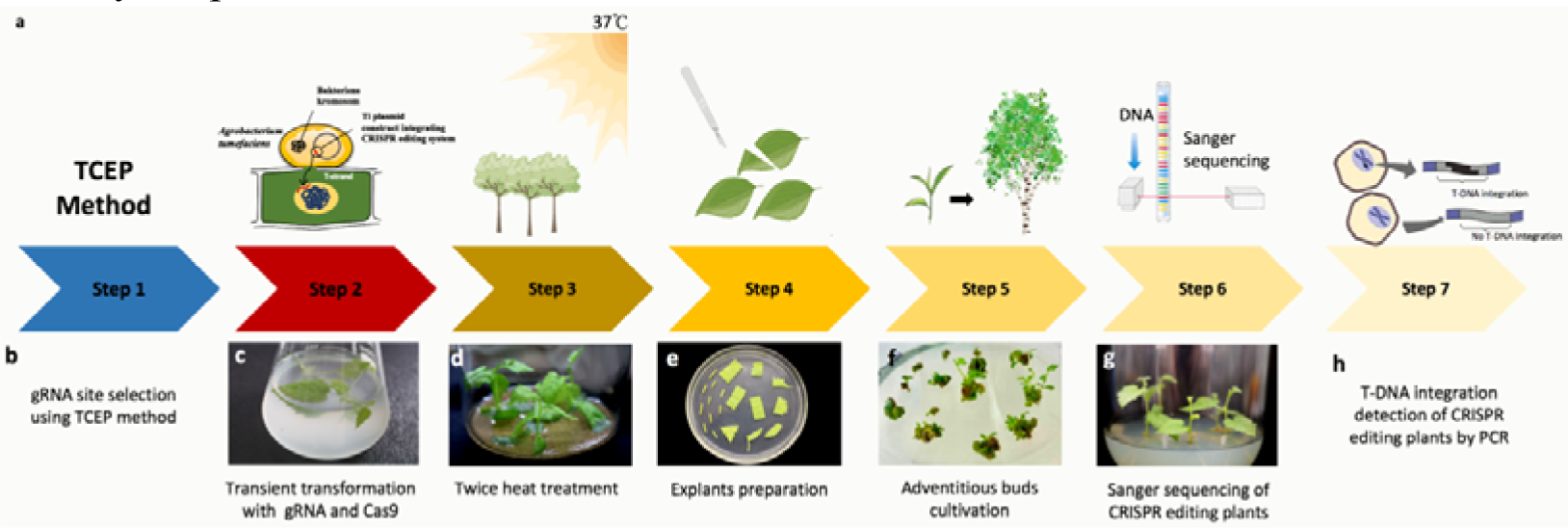
Flow chart of the CPDAT procedure. The procedure of CPDAT includes gRNA site selection, transient transformation, heat treatment to increase Cas9 activity, explants differentiation and growth, CRISPR-edited plants detection, and selection of CRISPR-edited plants without D NA integration.

### Selection of target editing site of *BpEBP1*

In the present study, we used the *BpEBP1* gene from Birch for the CRISPR editing study. First, we identified the ideal target site that Cas9 can effectively cleave. In total, 4 target sites (Fig. 3a) on *BpEBP1* were designed and the cor responding sequence was cloned into pYLCRISPR/Cas9P35S-N vector respectively, and the cleaving activities of Cas9 to these sites were determined by TCE P method. Cas9 was chosen for further research after the qPCR findings revealed that it had the highest cleaving activity in the *BpEBP1-sg2* region (Fig. 3b).

**Fig 3.**
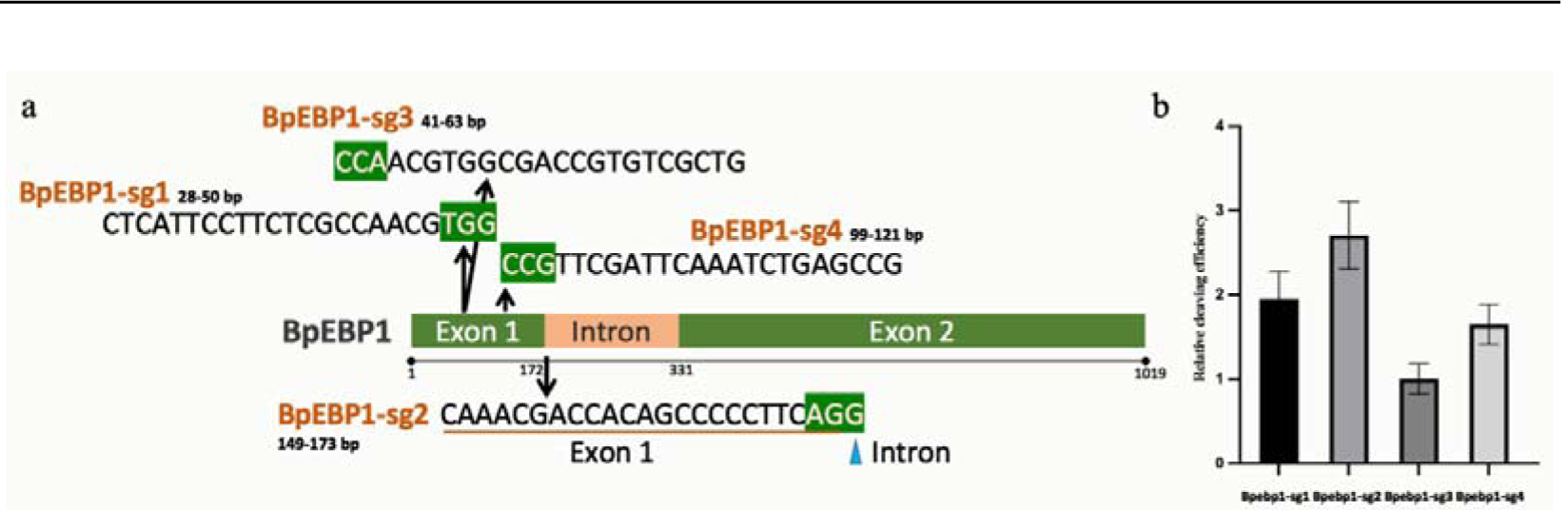
Location of gRNA sites and cleavage efficiency of these target sites in *BpEBP1*. a. The sequence and location of four target sites of *BpEBP1*. b. Determination of the cleaving efficiencies of Cas9 to the different target sites in *BpEBP1* using the TCEP method. The site with the lowest efficiency was set as 1 to calc ulate the relative cleavage efficiency.

### Generation of CRISPR-edited birch plants using CPDAT

We had developed a temporary transformation technique in the past (Ji et al., 2014; Zang et al., 2017). We also investigated the impact of sonication on the effectiveness of the transient transformation. The results indicated that sonication treatment (24w, 46KHz) can improve the efficiency of transient transformati on significantly, but long-time sonication also reduced the transformation efficie ncy. Especially, sonication with 15 s can increase the efficiency of transient transformation to a peak (Fig. 4a). The optimal density of Agrobacterium cells harboring pYLCRISPR/Cas9P35S-N was also determined. The results showed that the efficiency of transient transformation reached a peak when Agrobacterium cells of 0.7 OD600 (Fig. 4b). Therefore, for transient transformation, the optim ized conditions are that birch plantlets are treated with sonication (24w, 46KHz) for 15 s, and the Agrobacterium OD600 is 0.7.

**Fig 4.**
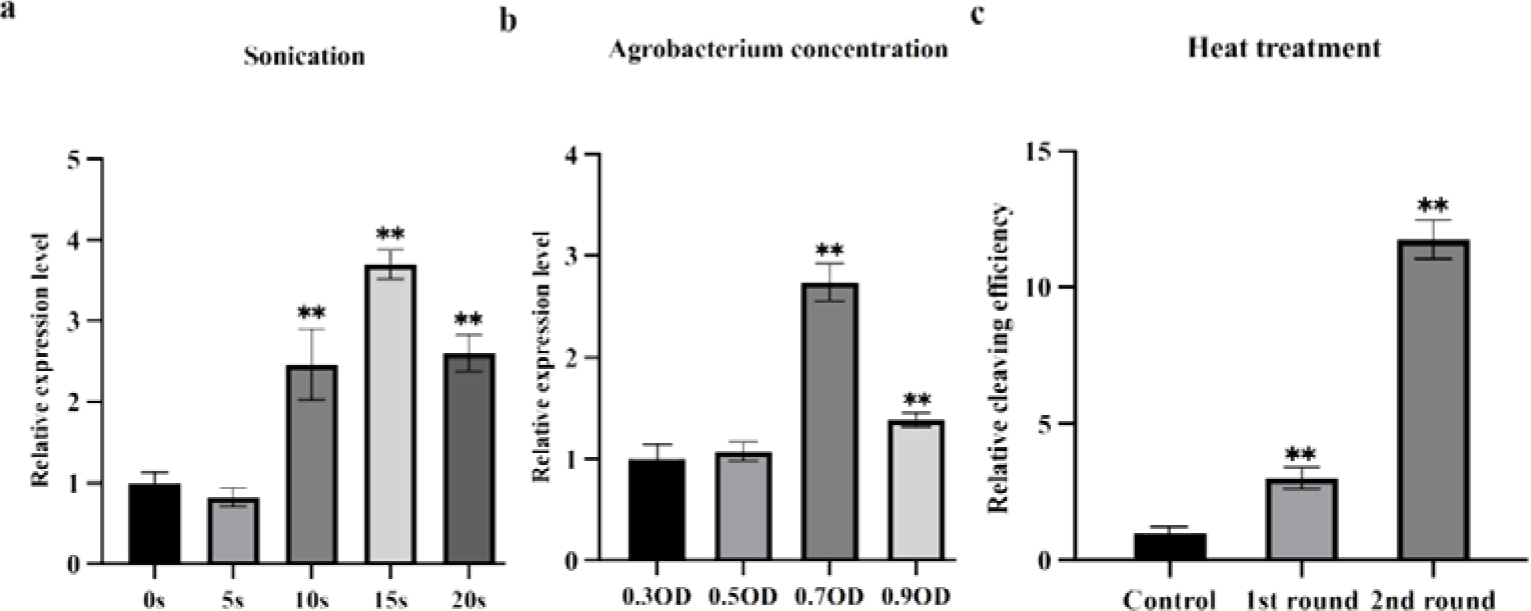
Optimization of the transient transformation and heat treatment on Cas9 activity. a: the effect of sonication on transient transformation efficiency. The birch plantlets were treated with sonication before the transient transformation. b: The eff ect of Agrobacterium density on transient transformation efficiency. The different cell densities were used for transient transformation. When determining the relative expression level, the Agrobacterium concentration with 0.3OD was used as the starting point. c: Determination of heat treatment (37°C) on Cas9 cleaving activity after transient transformation. After the transient transformation for 48 h, the birch plants were incubated at 37°C for 24 h, and incubated at nor mal conditions for 24 h, then treated at 37°C for 24 h again. After treatments, some birch plants were harvested immediately for TCEP detection. Significant differences were determined using Student’s t-tests: *P < 0.05, **P < 0.01. Heat treatments were applied twice to increase Cas9’s cleavage effectiveness. Following the TCEP approach, qPCR was carried out to ascertain whether heat treatment can increase cleavage efficiency. The results showed that heat treatment can significantly improve cleavage efficiency. Especially, the second round o f heat treatments can more significantly improve the cleavage efficiency (Fig. 4 c). These results indicated that heat treatment can significantly increase the cleavage efficiency of Cas9. Therefore, heat treatment is an important step in this procedure. After recovery for 7 d, the transiently transformed birch plants wer e cut into explants and grew in a culture medium to induce shoots without antibiotic selection pressure (Fig. 2e and 2f). When the shoots were induced, they were moved to a rooting medium for strength growth and generating roots (Fig. 2). These plantlets were potential CRIPSR-edited plants.

### Detection of the CRISPR-edited birch plant lines using Cruiser^TM^ digestion

We randomly selected 65 independent lines for CRISPR editing analysis. Cruis er^TM^ digestion was first performed to determine whether these lines had been mutated by CRISPR editing. The PCR product containing the target site with 6 60 bp in length from each line and WT plants were mixed. After the mixture, the PCR products were denatured and then renatured, then were digested with Cruiser^TM^, and the digested product was analyzed using agarose gel electrophoresis. The findings revealed 52 lines with two bands, indicating that these 52 lines have undergone mutations (Fig. 5). Therefore, the mutation rate is 80.00% (52/65). The remaining 13 lines with one band were not mutated by CRISPR.

**Fig 5.**
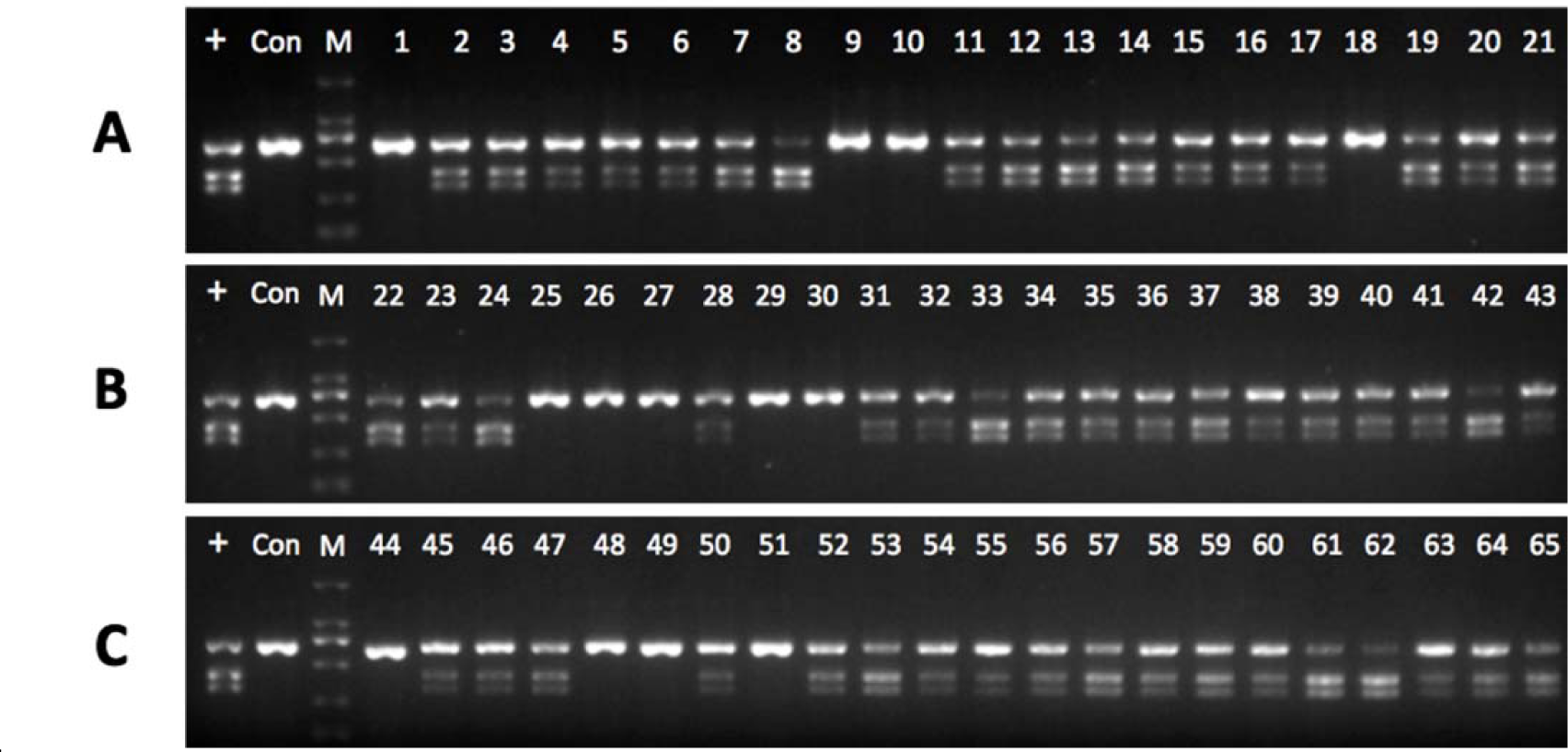
Cruiser^TM^ enzyme digestion assay of 65 randomly selected lines. The primers were designed to amplify the DNA region containing the gRNA site with 660 bp in length. The PCR product from each line and WT plant wer e mixed, denatured, then renatured, and digested with Cruiser^TM^ enzyme. The d igested products were analyzed by agarose gel electrophoresis. (+), The PCR from a previously identified CRISPR-edited plant and WT birch plants were mixed and was used as a positive control; (-), a digestion reaction with PCR prod uct of WT birch plants was as a negative control; M, DL2000 marker.

### Determination of the mutation of *BpEBP1* by Sanger sequencing

For a better understanding of the target situation, the PCR product containing t he target site of 65 lines was constructed into a pMD^TM^18-T vector for Sanger sequencing. Five clones from each line were randomly selected and sequenced, and a total of 325 clones were sequenced. Following sequencing, we discovered that, following the Cruiser^TM^ digestion, 52 of the lines had undergone CRISPR editing and 13 did not. Among them, 40.00% of lines with both alleles mutated, including 23.08% of homoz (A_1_/A_1_), 13.85% of biallelic (A_1_/A_2_), and 3. 08% biallelic/chimera (A_1_/A_2_/A_3_, i.e., containing two types of mutation forming chimera). The remaining 60.00% includes 20% of no mutation and 40% heterozygous and/or chimera (Table 1).

**Table 1.** Rates of mutation types.

| Lines with both alleles altered or chimera<br>(26 lines/40%) |  |  | Heterozygous or chimera<br>(39 lines/60%) |  |  | Mutation<br>rate |
| --- | --- | --- | --- | --- | --- | --- |
| Homozygous<br>($A_1/A_1$) | Biallelic<br>($A_1/A_2$) | Biallelic Chimera<br>($A_1/A_2/A_3$) | Heterozygous or<br>Chimera ($A_1/W$) | WT<br>($W/W$) | Chimera/chimera<br>($A_1/A_2/W$) | |
| 15/65<br>(23.08%) | 9/65<br>(13.85%) | 2/65<br>(3.08%) | 25/65<br>(38.46%) | 13/65<br>(20.00%) | 1/65<br>(1.54%) | 52/65<br>(80.00%) |

Note: A1, A2, and A3: the different mutated sequences; WT: The wild-type sequence (no mutation).

### Generation of CRISPR-edited birch plants without DNA integration

Since this technique used a transient transformation, some CRISPR-edited lines —those that lack CRISPR-edited DNA—might not integrate with T-DNA. To determine the CRISPR-edited lines without DNA integration, PCR detection was performed to determine T-DNA insertion in different lines. The T-DNA region s including the left, middle, and right regions of T-DNA were PCR amplified, respectively. The results showed that there were 5 lines out of the 52 mutated lines were not integrated with T-DNA, indicating that they were CRISPR-edite d lines without DNA integration (Fig. 8). Therefore, the rate for generating CR ISPR-edited lines without DNA integration was 7.69% (5 from 65 checked line s). For further determining the mutation types of these CRISPR-edited plants w ithout DNA integration, the results of Fig. 6 and Fig. 7 were combined for an alysis. The results showed that these mutations include 2 homozygous and 3 bi allelic, but no heterozygous/chimera type was found. (Table 3).

**Figure 6.**
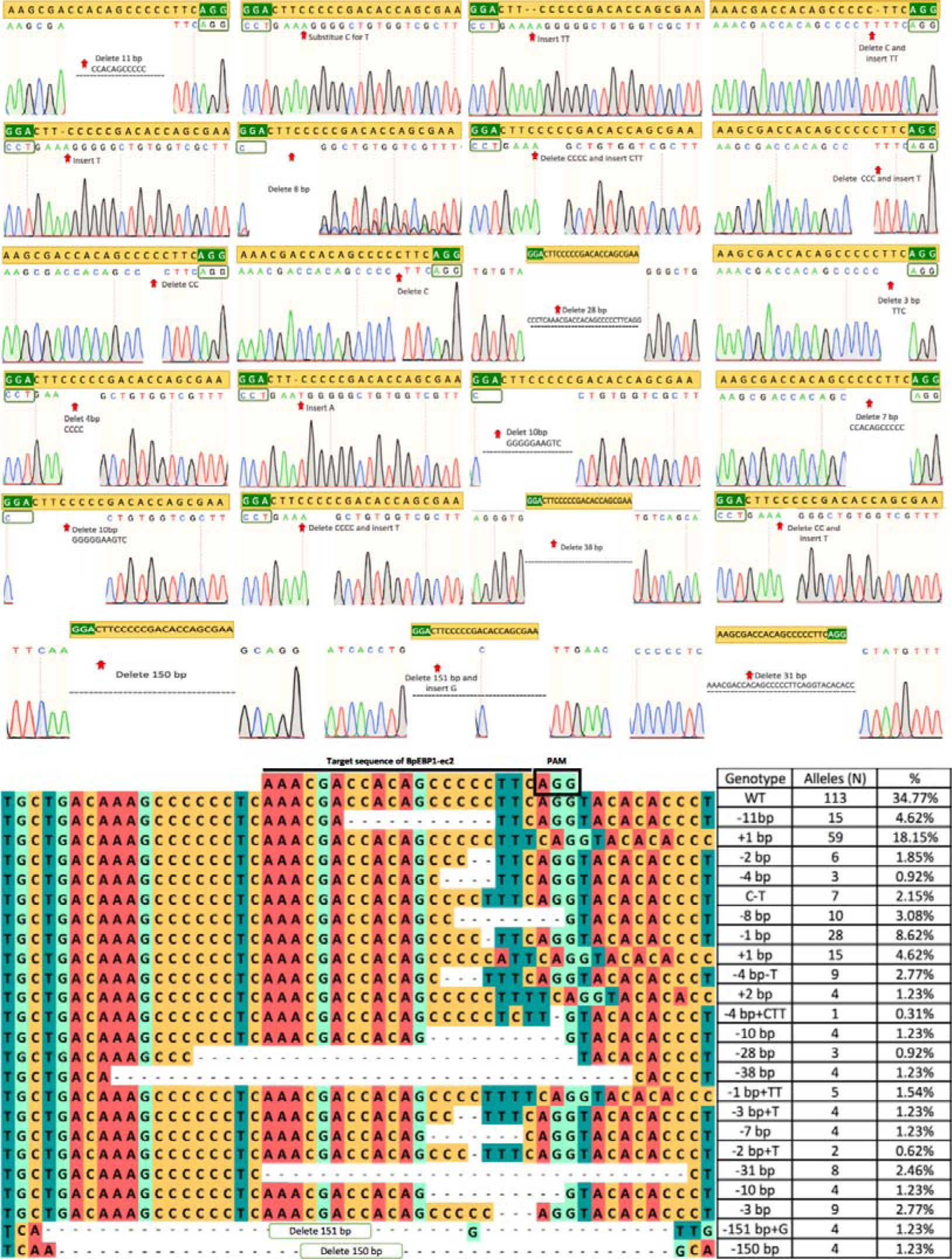
Genotypes of 23 representative mutation birch plants identified b y Sanger sequencing. In total, 65 lines were selected and the target regions w ere PCR amplified. The PCR product was cloned into a pMD^TM^18-T vector fo r Sanger sequencing. Five clones from each line were selected for sequencing, and 325 clones in total were sequenced. In this chart, each clone can be consi dered an allele. Allele: the sequence from a clone. WT: the sequence without mutation. +, insertion; -, deletion; -A, A bp deletion; +B, B insertion; a-b, b substitute for a. Out of 325 clones, 113 were found to be mutant-free, while the remaining 212 displayed 23 different types of mutation events, according to the sequencing results (Fig. 6). Among the 23 types of mutation events, 83 lines were mutated by deletion of 15 bp or less, and 19 lines had large deletion, 78 lines were mutated by small insertion (15 bp insertion or less), 28 lines were mutated by small substitution (15bp substitution or less), and 4 lines were mutated by large substitution (more than 15bp substitution) (Table 2 and Fig. 6).

**Fig 7.**
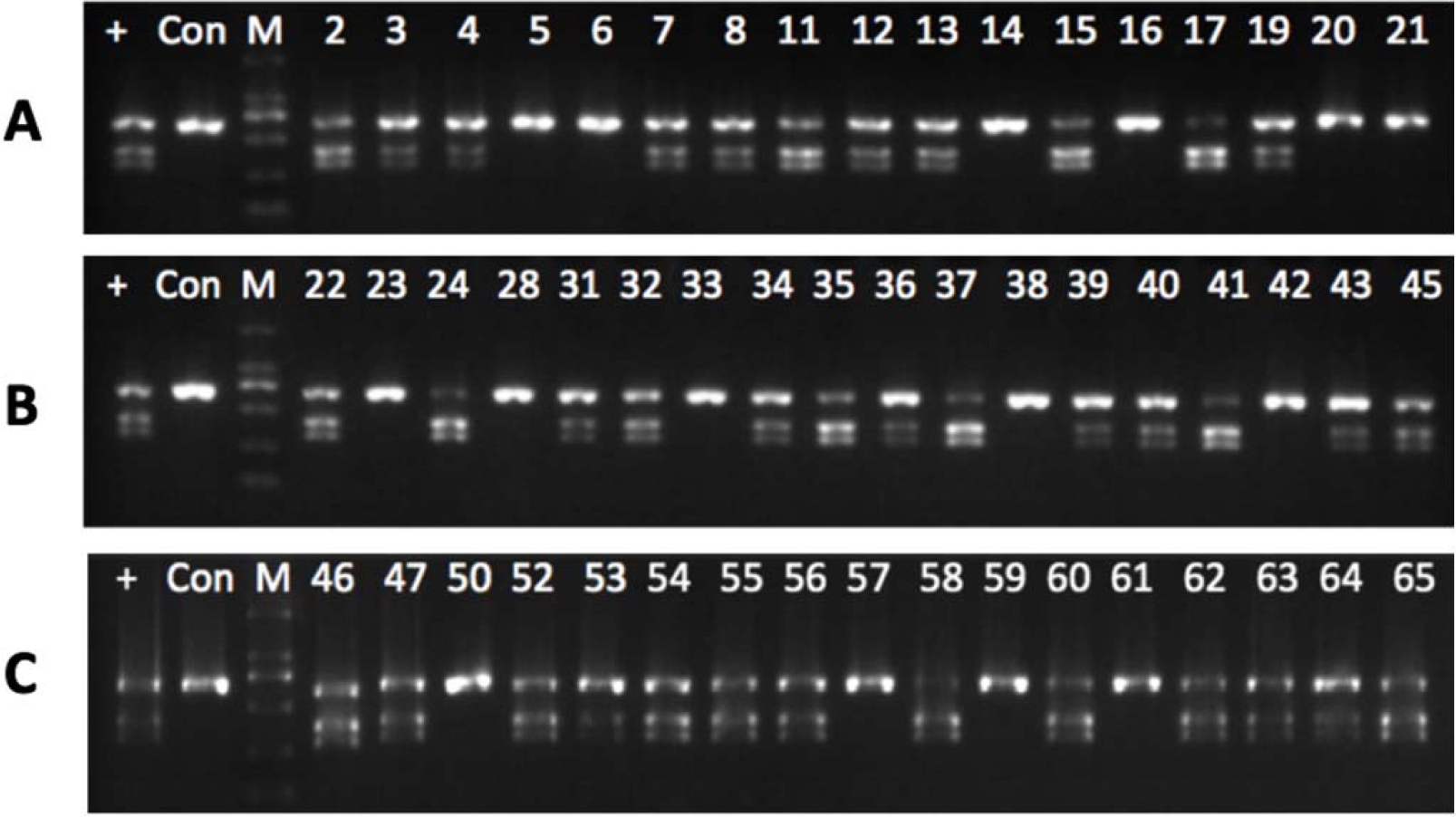
Determination of homozygous, heterologous mutation, or heterologous/chimera. The PCR product from each line was denatured, renatured, and then digested with CruiserTM. The digestion product was analyzed using agarose gel electrophoresis. (+), The PCR from a previously identified biallelic CRISPR-edited birch plant was used as a positive control; Con, a digestion reaction with PCR product of WT birch plant; M, DL2000 marker.

**Figure 8.**
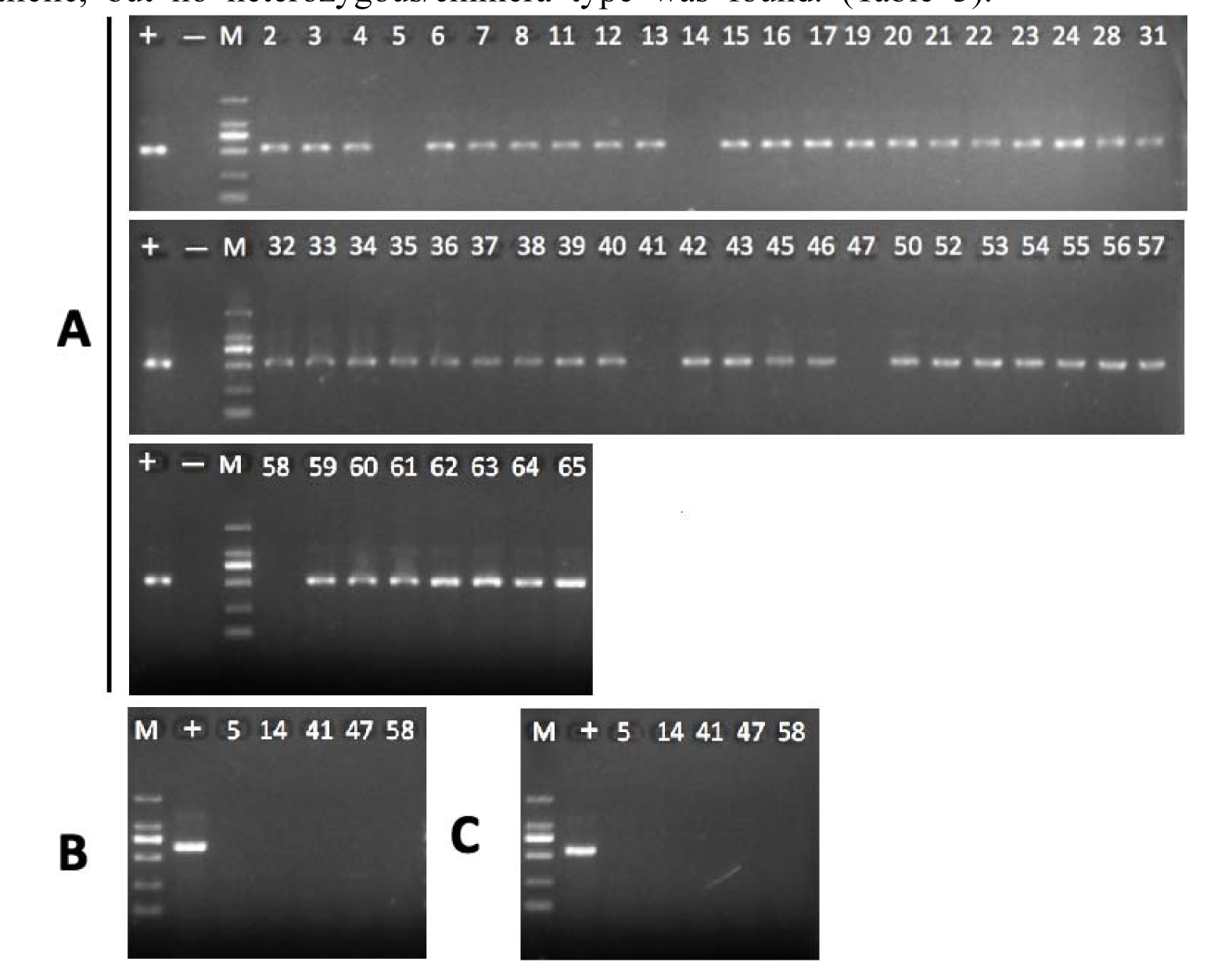
Detection of CRISPR-edited birch lines without DNA integration. All the selected birch plants were PCR amplified to detect whether T-DNA ha d been integrated. The detected PCR amplification regions were the left, middle, and right regions of T-DNA. (-), The product from WT was as a negative control; (+), the PCR product from an identified transgenic line was used as the control.

**Table 2.**
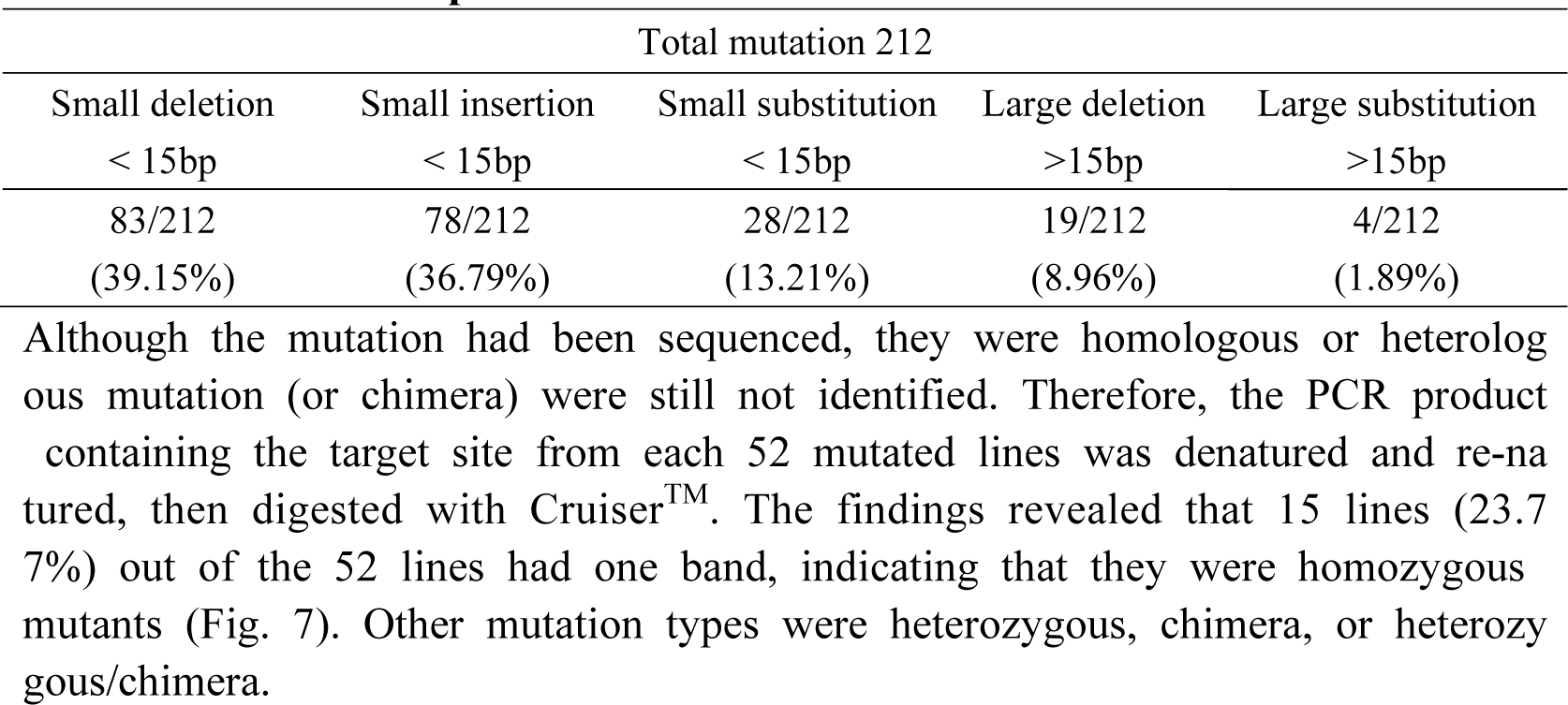
The mutation patterns.

| Total mutation 212 |  |  |  |  |
| --- | --- | --- | --- | --- |
| Small deletion<br>< 15bp | Small insertion<br>< 15bp | Small substitution<br>< 15bp | Large deletion<br>>15bp | Large substitution<br>>15bp |
| 83/212 | 78/212 | 28/212 | 19/212 | 4/212 |
| (39.15%) | (36.79%) | (13.21%) | (8.96%) | (1.89%) |

**Table 3.**
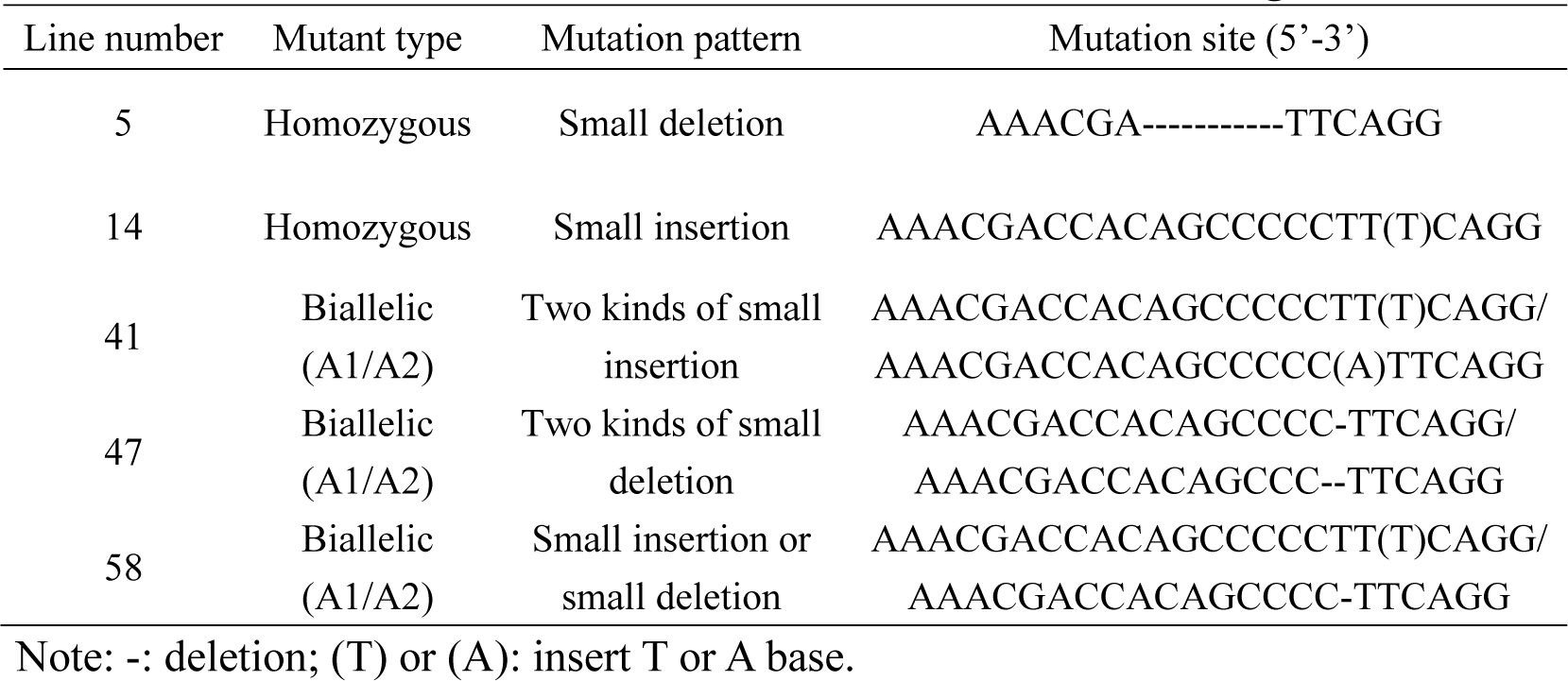
Mutation of CRISPR-edited birch lines without DNA integration.

## Discussion

### Enhancing Efficiency Through the CPDAT Method for Generating CRISP R-Edited Birch Plant

In this method, the birch plants were transiently transfor med to cleave the target site for inducing mutation. The adventitious buds wer e produced after transient transformation without antibiotic selection pressure, which may have two implications (Figs. 1 and 2). One is that the differentiation rate of adventitious buds without antibiotic selection pressure is as similar as under normal conditions, and many adventitious buds will be generated. The other is that the T-DNA may not be integrated into the genome without antibiotic selection pressure. Therefore, the CRISPR-edited birch plants without DNA integration may be generated, which could be identified by selecting the lines without T-DNA integration from the CRISPR-edited plants. In the present study, many adventitious buds have been generated. Furthermore, about 80% of the lines that were produced were CRISPR-edited, indicating that this technique could significantly increase the efficiency of producing CRISPR-edited plants (Fi g. 5). Moreover, our results showed that 7.69% of the generated plants were n ot integrated with any T-DNA (Fig. 8), and these mutations were homozygous and biallelic (Table 3). These results indicated that CRISPR-edited birch plants without DNA integration can be generated with a high percentage. What dese rved to be mentioned is that most of the mutations (89.15%) are small substitutions, small deletions, or small insertions (Table 2), suggesting that the error-prone mechanism NHEJ plays an important role in repairing DSBs.

### Factors affecting the generation of CRISPR-edited birch plants

The most important factor is the high efficiency of transient transformation which is quite important for generating CRISPR-edited birch plants. In this study, we first optimized the efficiency of the transient transformation of birch. Also, the birch plants should be in a good state after transient transformation, which is quite important for explant differentiation. The second factor is that heat treatment is used to enhance the cleaving activity of Cas9, which will induce more DNA cleaving resulting in more mutation, and the mutation can be accumulated in CRISPR editing. Previous studies showed that heat stress (37°C) can significantly increase targeted mutagenesis by CRISPR/Cas9 in birch plants(LeBlanc et al., 2018). In the present study, we used heat treatment to increase t he cleaving activity of Cas9, which could make more genomes to be cleaved. As the mutation in the genome can be accumulated, two rounds of heat treatment are necessary, which could accumulate more mutation (Fig. 4c). The birch plant should have a tissue culture regeneration mechanism, which is the third factor. Because the birch plants were differentiated without antibiotic selection pressure, this system does not require a high frequency of regeneration, but the regeneration mechanism is required to produce the adventitious buds. (Fig. 2).

### Method for avoiding chimera lines in CRISPR-edited birch plants

In the present study, the CRISPR-edited birch plants without DNA integration can be selected directly by PCR detection. However, despite the absence of antibiotic selection pressure, our data also demonstrated that T-DNA was integrate d into the genome to create transgenic lines with a very high percentage (Fig. 5). This may be due to the following reasons. One is a high percentage of cells had been transiently transformed, and a high level of T-DNA in the nucleus makes T-DNA integrated into the genome with a high frequency. The other reason is that some T-DNA integrated lines may be a chimera, i.e., the CRISPR-edited plants contain WT cells. According to Cruiser^TM^ and DNA Sanger sequencing analysis, heterozygous or chimera accounted for 60.00% (Table 1). Among the heterozygous or chimera lines, there is maybe a high percentage of chimera. However, Cruiser^TM^ digestion and DNA Sanger sequencing analysis cannot distinguish the heterologous and chimera plants. For CRISPR-edited lines without DNA integration, to get rid of the chimera lines, the method of Cruis er^TM^ digestion can be used to directly select the homozygote and biallelic lines. For this reason, the Cruiser^TM^ digestion method could directly detect homozyg ous mutation (Fig. 7), and Sanger sequencing with more clones to be sequence d can identify the biallelic mutation (Fig. 6).

For the common CRIPSR-edited lines (i.e., not CRISPR-edited lines without D NA integration), in addition to the method by directly selecting homozygote an d biallelic lines using Cruiser^TM^ and DNA sequencing method, the other method is that the CRISPR-edited birch plants can be cut into explants to differentiate under antibiotic selection pressure. For the reason that most of the generated CRISPR-edited lines are integrated with T-DNA in the genome and harbor th e antibiotic gene, the differentiation of adventitious buds will not take a long t ime and will have a high rate. Each adventitious bud can be considered as an independent line and analyzed using Cruiser^TM^ and DNA Sanger sequencing t o select the CRISPR-edited line. The adventitious buds containing chimeras wil l be significantly fewer because they are developed from just one or a few cells.

## Conclusion

In this present study, we provided a novel strategy and method for the generation of CRISPR-edited birch plants without DNA integration. This technique exhibits improved efficiency for producing conventional CRISPR-edited birch plan ts in addition to its ability to generate CRISPR-edited birch plants without DN A integration. This method could adapt to the plant that can be infected by *A. tumefaciens* and has a tissue culture regeneration system. Importantly, this met hod offers a streamlined approach to generating CRISPR-edited birch plants mo re easily than generating transgene plants. This innovation holds significant pot ential for wide applications in plant molecular breeding techniques.

## MATERIALS AND METHODS

### Plant material and growth conditions

The birch plantlets grew in a culture medium and were used for study. For ad ventitious bud induction, the adventitious bud induction medium (ABI) was use d (Wood Plant medium [WPM] + 1 mg·L^-1^ 6-BA, pH 5.8) to generate callus and induce adventitious bud simultaneously. For root generation and plantlet gr owth, a rooting medium (RM) was used (WPM + 3% (w/v) sucrose + 0.2 mg /L NAA) and used for transient transformation. The plantlets were cultured und er conditions with 16 h/8 h of light/dark cycle, 70% relative humidity, light in tensity 40 μmol s^-1^ m^-2,^ and temperature of 24°C.

### Target design and construction of CRISPR/Cas vector

The gene for *BpEBP1* (Genbank number: MW202047.1) from birch was used f or the study. All protospacer adjacent motifs (NGG) in *BpEBP1* were screened by the online program (http://skl.scau.edu.cn), and 4 target sites were selected for the study. The primer pairs for gRNA were cloned into the pYLCRISPR/ Cas9P35S-N vector using the golden gate assembly method to construct a paire d-sgRNA/Cas9 binary. The constructs were transferred into *A. tumefaciens* (strai n EHA105) by electroporation method. To determine the target site with high Cas9 cleavage efficiency, the TCEP method was employed. The procedure of t he TCEP method and algorithm for calculation of the CRISPR cleavage efficie ncy were following the description of Wang et al(Wang et al., 2021).

### Transient genetic transformation

The transient genetic transformation was performed according to Zang et al.(Za ng et al., 2017) with some modifications. In brief, colonies of *A. tumefaciens* were cultured in an LB liquid medium supplied with 50 mg/L rifampicin and 50 mg/L kanamycin at 180 rpm at 28°C. After the cell density OD600 to 1.0, 1 mL of culture was transferred to 50 mL of fresh liquid media and was cul tured to an OD600 of 0.7. The cells were then collected by centrifugation at 2 000×g for 10 min, and were adjusted to an OD600 of 0.8 with a solution [1/2 MS+2 mM MES-KOH +200 mg/L DTT (Dithiothreitol)+20 μM 5-Azacytidine +2.5% (w/v) sucrose +10 mM CaCl_2_ +120 μM AS (Acetosyringone) +270 m M mannitol +Tween 20 (0.01%, v/v), pH 5.8], and was served as transformati on solution. For transient genetic transformation, the whole birch plants were s oaked into 200 ml of transformation solution at 100 rpm for 2 h at 25 °C. Th e soaked birch plants were moved to a solution (1/2 MS +200 mg/L DTT +10 0 μM AS +2.5% (w/v) sucrose +270 mM mannitol) washed for 30 seconds an d then wiped with sterile filter paper to remove the excessive *A. tumefaciens* cells. The birch plantlets were planted vertically on 1/2 MS solid medium [1/2 MS +1% (w/v) sucrose +200 mg/L DTT+ 100 μM AS, pH 5.8] for 48 h. The transformed birch plants were treated at 37°C for 24 h (harvested some mater ial as sample 1 immediately after treatment), recovered at 25°C for 24 h, then treated at 37°C for 24 h (harvested some material as sample 2 immediately after treatment) again. After treatments, the birch plants grew at the normal condition for 7 d, and these transiently transformed plants were cut into explants and cultured in an ABI medium for inducing buds (ABI +300 mg/L cefotaxime sodium +1 mg/L AgNO_3_). Each bud can be considered as an individual lin e since when the buds were induced, RM media (RM + 300 mg/L cefotaxime sodium) was employed for growth and the generation of roots.

### Identification of mutation through T7E1 digestion and Sanger sequencing

Total DNA was extracted from birch using a one-step method for rapid extract ion of birch plant genomic DNA reagents (BioTeke Corporation, Wuxi, China). The PCR-paired primers were designed to amplify the DNA region containing the gRNA target site with 567 bp in length. The paired primers were: EBP1- F (5’-CCCTCGGGTTTGGCTGGATGAC-3’) and EBP1-R (5’-ACCTTTCTCGG GTCGCGTA-3’). The PCR reaction system is as follows: 1 μL DNA (0.1-0.5 μg DNA), 2 μL 10× PCR buffer, 0.5 μL EBP1-F/EBP1-R (10μM), 0.2 μL Ex Taq (Takara Biomedical Technology, Beijing, China), 0.4 μL dNTP (10 mM each) with the total volume of 20 μL. The PCR thermal profile is set as 94°C 3 min, 30 cycles of 94°C 15 s, 58°C 30 s, and 72°C 60 s. The PCR product was purified with a PCR purification kit (Takara Biomedical Technology, Bei jing, China) and was cloned into a pMD^TM^18-T vector (Takara Biomedical Technology, Beijing, China). The construct was transformed into DH5α *Escherichia coli*, and 5 clones were randomly selected from each construct for Sanger seq uencing.

### Cruiser^TM^ enzyme digestion assay

**Cruiser^TM^ enzyme**(Genloci Biotech, Nanjing, China) was used to detect mutations induced by CRISPR editing. The PCR product was amplified from the total DNA of the potentially CRISPR-edited plants (transgenic line) or WT birch plants using the primers of EBP1-F and EBP1-R. A reaction system contains 20 0-400 ng PCR purified product and 1 μL of 10× Cruiser^TM^ reaction buffer and was denatured at 95 °C for 3 min, then incubated at 68 °C for 2 h, added with 1 μL of Cruiser^TM^ at 37 °C, and incubated at this temperature for 30 mi n. For determining whether CRISPR editing occurred, PCR products from the transgenic line and WT were mixed. The PCR output from a single transgenic line was utilized to identify homologous, heterologous mutations, or chimeras. Electrophoresis on 2% agarose gel was used to examine the Cruiser^TM^ digestio n product.

### Determination of CRISPR-edited lines without DNA integration

The CRISPR-edited birch plants were further screened to determine whether the y were CRISPR-edited plants without DNA integration or not. Three paired pri mers were respectively designed to amplify the left, middle and right regions o f T-DNA. The PCR was performed using the template of total DNA from the CRISPR-edited lines. The CRISPR/Cas9 plasmid was used as a positive control, and ddH_2_O was as a template to serve as the negative control. The primers used were shown in Supplementary Table S1.

### Real-time PCR

For determining the cleavage efficiency of Cas9 in different gRNA sites and b y heat treatment, real-time PCR was performed. Total DNA was isolated from the birch using a one-step method for rapid extraction of birch plant genomic DNA reagents (BioTeke Corporation, Wuxi, China). The real-time PCR reaction system is as the followings: 10 µL of SYBR Premix Ex Taq™ (Takara Biom edical Technology, Beijing, China), 0.5 µM of each forward or reverses primer and 100-500 ng of DNA template. Real-time PCR was performed on qTower 2.2 real-time PCR machine (Analytik Jena AG, Jena, Germany). The thermal profile was 94 °C for 30 s; 45 cycles of 94 °C for 15 s, 58 °C for 30 s, and 72 °C for 45 s. The primers used were shown in Supplementary Table S2. T he figures were computed in accordance with Wang et al^30^. There were three b iological replications.

## Author contribution

YW conceived and directed the research. SS and XH performed the experiments and the overall data analysis; RJ and SN prepared figures and tables. DW, ZY and JW provided birch plants. SS and YW wrote the manuscript, all authors revised the manuscript. All authors read and approved the final manuscript.

## Funding information

This work was supported by Funds for the Xingliao Talent Plan Project XLYC1902007 and Guiding Local Scientific and Technological Development by the

Central Government 2020JH6/10500071.

## COMPETING INTERESTS

The authors declare that they have no competing interests.

## Data Availability Statements

The data underlying this article will be shared on reasonable request to the corresponding author.

